# A database resource for Genome-wide dynamics analysis of Coronaviruses on a historical and global scale

**DOI:** 10.1101/2020.02.05.920009

**Authors:** Zhenglin Zhu, Kaiwen Meng, Geng Meng

**Author notes:** Corresponding authors, Zhenglin Zhu, School of Life Sciences, Chongqing University, No.55 Daxuecheng South Road, Shapingba, Chongqing, 401331, China., Geng Meng, College of Veterinary Medicine, China Agricultural University, Beijing, 100094 China.

## Abstract

The recent outbreak of a new zoonotic origin Coronavirus has ring the bell for the potential spread of epidemic Coronavirus crossing the species. With the urgent needs to assist the control of the Coronavirus spread and to provide valuable scientific information, we developed a coronavirus database (CoVdb), an online genomics and proteomics analysis platform. Based on public available coronavirus genomic information, the database annotates the genome of every strain and identifies 780 possible ORFs of all strains available in Genebank. In addition, the comprehensive evaluation of all the published genomes of Coronavirus strains, including population genetics analysis, functional analysis and structural analysis on a historical and global scale were presented in the CoVdb. In the database, the researcher can easily obtain the basic information of a Coronavirus gene with the distribution of the gene among strains, conserved or high mutation regions, possible subcellular location and topology of the gene. Moreover, sliding windows for population genetics analysis results is provided, thereby facilitating genetics and evolutional analysis at the genomic level. CoVdb can be accessed freely at http://covdb.popgenetics.net.

## Introduction

Coronavidae is a group of positive-sense, single-strand RNA viruses with a likely ancient origin, and human Coronavirus repeatedly emerged during the past handred years^1^. Coronaviruses are classified into four distinct genera: alpha and beta Coronavirus mainly infect mammals, whereas gamma and delta Coronavirus more circulate in avian hosts^2^. As a potential dangerous zoonotic disease, the previous outbreaks of respiratory syndrome-related Coronavirus (SARS-CoV) and Middle East respiratory syndrome-related Coronavirus (MERS-CoV) have plagued the general public and researchers in the past years ^3^. Recently, a new Coronavirus (2019-nCoV), which may originated from wild animals, first identified in Wuhan city, China. Till now it has been resulted in more than fourteen thousand confirmed infections in China^4, 5^ with the cases number is still increasing. Although we have knowledge and experience in the virology, diagnosis, clinical characteristics, and other aspects related to SARS-CoV and MERS-CoV, there are many unanswered questions about the new emerging 2019-nCoV. The new Coronavirus outbreak in China strongly reminds the continued threat of zoonotic diseases caused by Coronavirus to global health security. Sharing experience and knowledge from across disciplines in the historical and global scale should provide valuable scientific knowledge to fight against the threat of Coronavirus.

The aim of the construction of CoVdb is to provide Coronavirus knowledge, to contribute to global Coronavirus research, especially for the investigation of the emerging 2019-nCoV. For previous works, ViPR^6^ and ViralZone^7^ are general data resources and is lack of analysis tools in population genomics and evolution. In contrast, CoVdb is specially designed for Coronavirus. It combines, compares, and annotates all the published Coronavirus genomes up to date^8,9,10,11,12,13,14,15,16,17,18,19^. The new developed database provides the convenience for the identification of gene function and identity among the Coronaviridae genomes. CoVdb provides information on subcellular location, functions, proteins topology, as well as population level thorough analysis results. We will be dedicated to keep updating all the genomic information and optimizing the database.

## Result and discussions

### Data and information

CoVdb extensively collects published Coronavirus genome data (Table S1), including 104 strains are of 780 possible ORFs. In average, there are 5-14 possible protein coding genes in each strain. Although the structure of Coronavirus (Figure 1) is not complex, we still performed a subcellular localization analysis of the Coronavirus genes to predict their roles in the infection process with addition to the previous research. Base on prediction only, 27% of the proteins are predicted to be located in the host nucleus or host cytoplasm (Figure S1). 32% of the Coronavirus genes are predict to be membrane proteins. In gene ontology, Coronavirus genes enrich in association to the membrane (Figure S2). Moreover, based on the population genetics test analysis, we found that 2242 regions are of a significantly low Tajima’s D^20^ and significantly high composite likelihood ratio, CLR^21, 22^ (Rank Test, P-value<0.05), indicating that these genes were recently possibly under positive selection (Table S2). These regions deposited in 400 genes, and most of the regions are located at ORF1 of the virus genome. Among all these 2242 regions, 98 are located in the noncoding region and 30 are located in the coding regions of non-structural proteins. We also found that most high CLR regions (1452, 64.8%) are at the protein ORF1ab (Table S3). These positive selection regions may involve in the adaptive mutations related to the virus replication and infection. The results are informative for future Coronavirus research and epidemiological survey.

**Figure 1.**
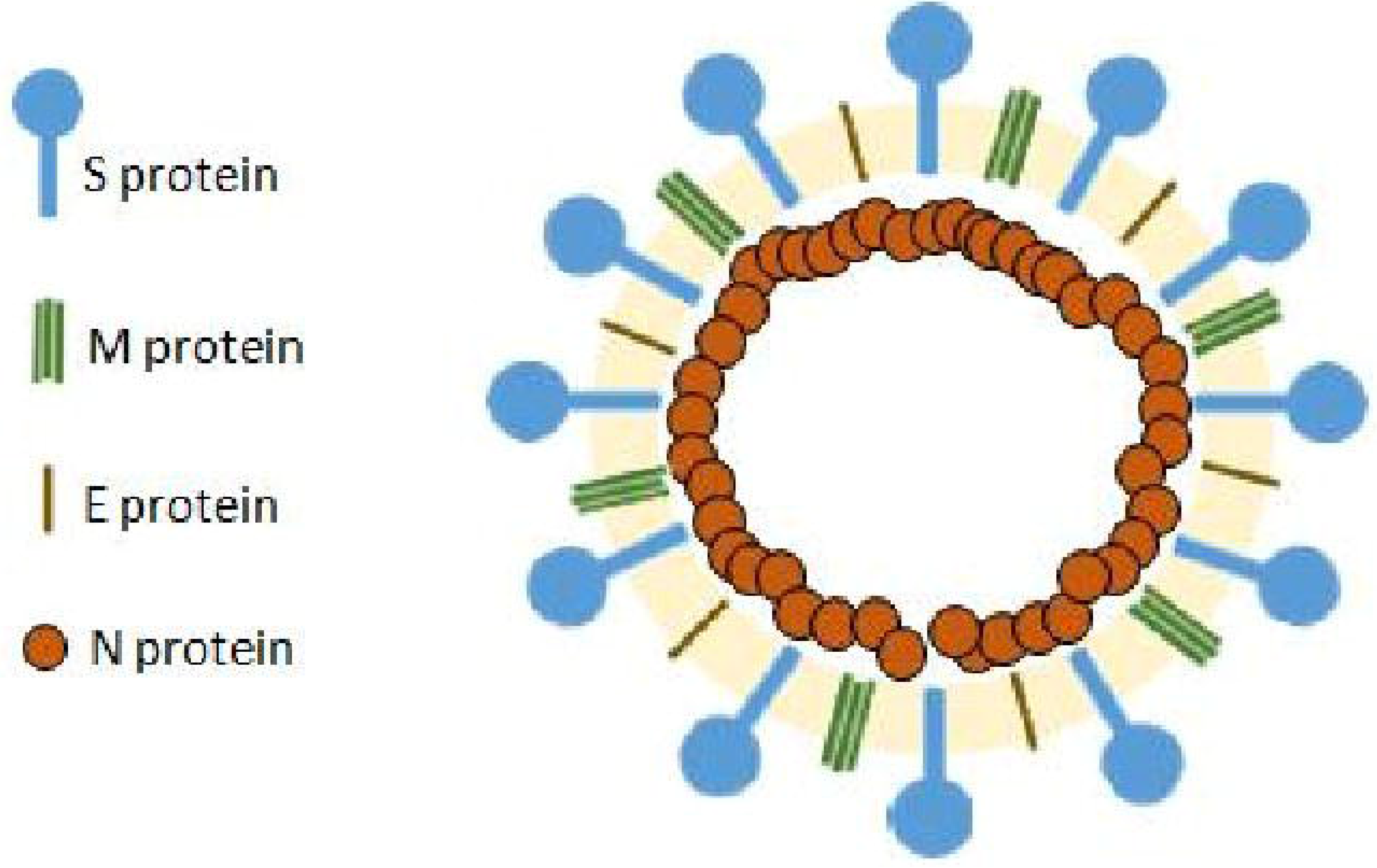
A diagram displaying the general structure of Coronavirus.

We identified 322 Coronavirus gene clusters from the genomes of 104 strains (for details, see Materials and Methods). Using the genomes documented in CoVdb, we generated the phylogenetic tree of Coronavirus (Figure 2). The tree indicates that 2019-nCoV is of close relationship with SARS-CoV and may arrive from bat. Coronavirus strains extracted from human are always surrounded by strains extracted from bats, indicating bats may be the main source leading to the infection of Coronavirus in human world (Figure S3).

**Figure 2.**
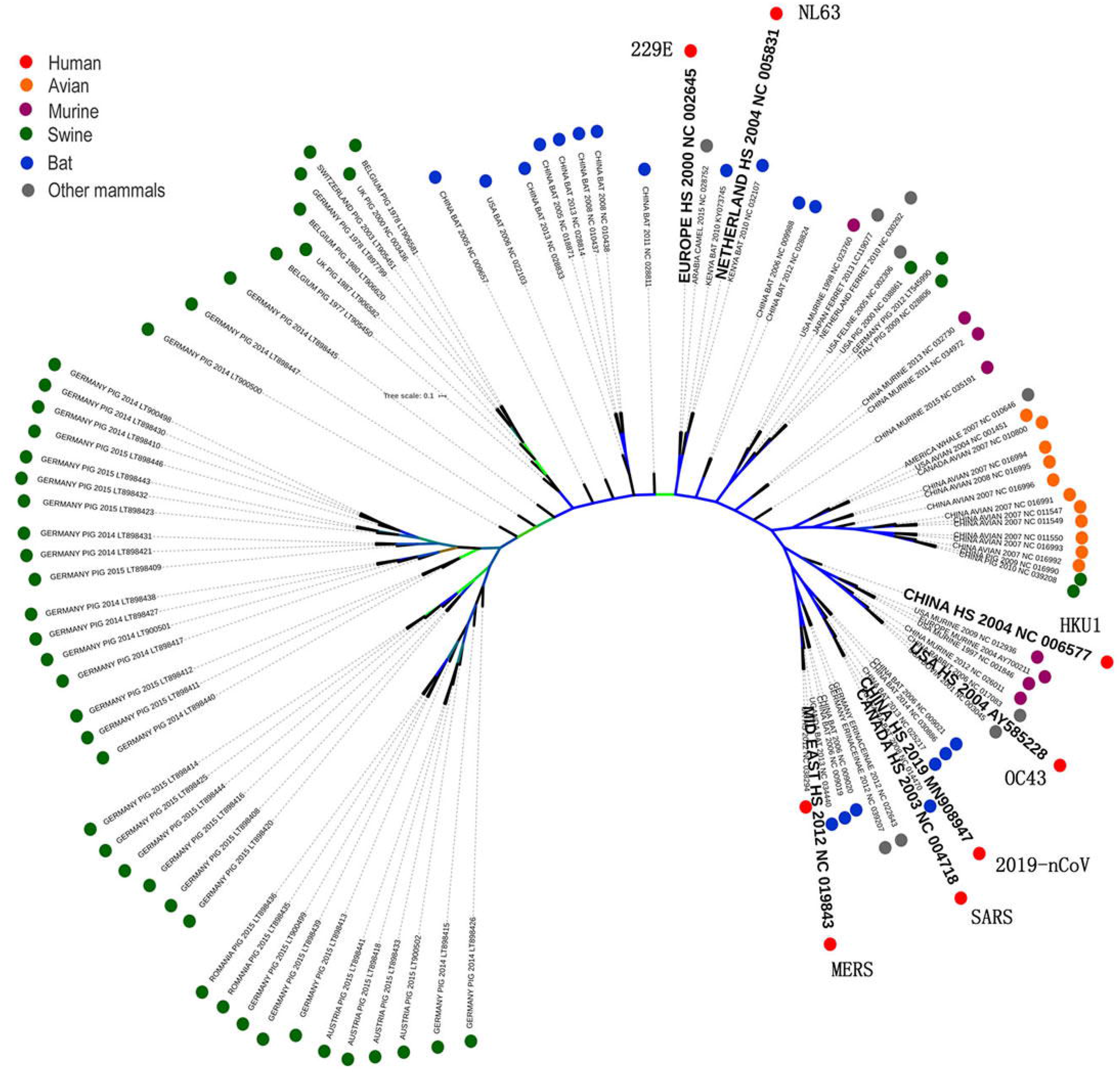
The phylogenetic tree built by Coronavirus genomes in CoVdb. Links with Bootstrap Likelihood Value = 1 are colored by blue and the ones with value > 0.5 is colored by green. The names of 7 major Coronavirus strains are enlarged. They are MERS, SARS, 2019-nCoV, OC43, HKU1, NL63 and 229E. Different hosts from which virus is collected are marked by cycles with different colors.

### Interface and main functions

The genome browser page follows a style with analysis tracks (Pi, Theta^23^, Tajima’s D^20^, and CLR^21, 22^) listed following gene segments (Figure S4). CoVdb are equipped with other general genome browser tools like UCSC genome browser^24^. In addition to basic information, CoVdb show gene information mainly in cell, protein structure and evolutionary signatures.

The search engine in CoVdb is powerful and supports fuzzy search, BLAT and Blast. CoVdb also allows to search by cell location, function, evolutionary test parameters and protein structure parameters (Figure 3). To facilitate personalized gene list analysis, CoVdb provides gene links through inputting a range of genomic locations or a list of genes’ accession numbers.

**Figure 3.**
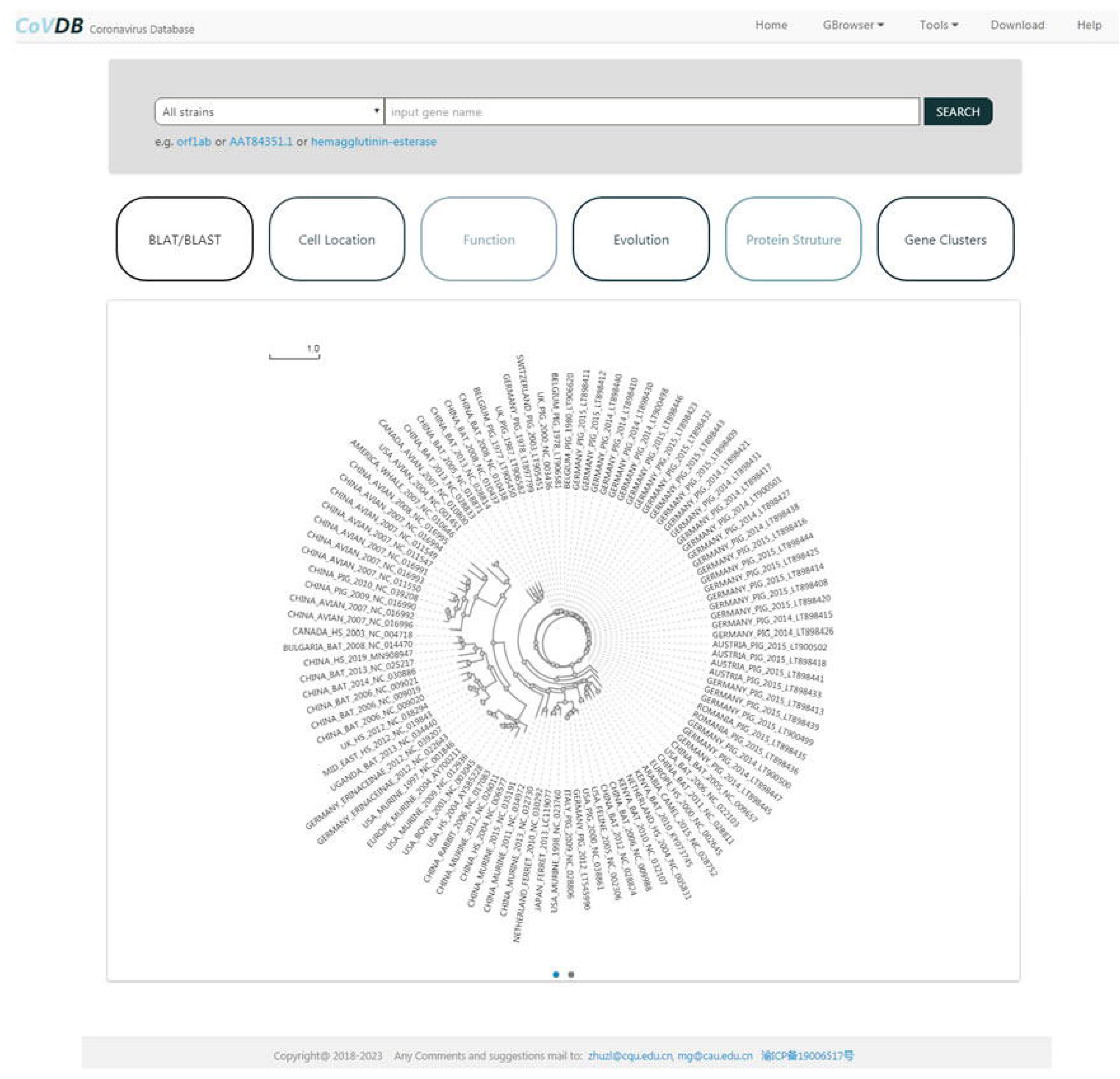
The main page of CoVdb.

## Conclusion

Dedicated to assist the researcher to combat the pandemic of 2019-nCoV, and to provide a more specialized platform for Coronavirus, we comprehensively gathered data and systematically constructed the Coronavirus Database, CoVdb. With the help of this database, we have successfully developed a novel tool (unpublished) to detect 2019-nCoV and the program is in the process to be put into production. Researchers can conveniently retrieve Coronavirus genomic and gene information from CoVdb. Hopefully, this database will play more important roles in fighting against the infection of Coronavirus.

## Materials and methods

### Gene Information, subcellular localization and topology prediction

Coronavirus sequences and annotations are downloaded from the NCBI genome database. For strains without ORF annotations, we reannotated the genomes through mapping known Coronavirus genes to the genome, requiring identity > 50% and coverage > 80%. Then, we verified the quality of these proteins by known proteins. We kept predicted proteins with both identity and coverage higher than 0.5. According to previous homologous gene identification methods^25, 26^, we also we performed pairwise alignments for all protein sequence, using CD-HIT^27^ with the parameters of identity > 90% and coverage > 70%. In this way Coronavirus proteins are clustered into 322 unified gene clusters.

We wrote PERL scripts to automatically BLAST Coronavirus protein sequence against the UniProt protein database^28^ and took the hits with the highest scores as the best matches requiring E-value <0.05. Using the accession number of matched UniProt proteins, we retrieved detailed proteomics information from UniProt. In the same way, we also acquired protein 3D structure information from PDB database^29, 30^.

We did the subcellular localization prediction of all Coronavirus genes using an online tool MSLVP^31^ We used TMHMM 2.0^32^ to predict the transmembrane helixes within protein sequences, and converted output images into PNG format by Magick (www.imagemagick.org).

### Evolutionary analysis

We utilized CUDA clustalW^33^ to perform a whole genome alignment of all Coronavirus genomes. The results are used to built a phylogenetic tree by FastTree 2.1^34^. We did genomic level alignment by LASTZ^35^, did sequence level alignment by MUSCLE^36, 37^ and made phylogenetic trees by FastTree 2.1^34^.

To detect selection signals, we did sliding widow analysis for each genome (window=2OO bp, step=50 bp) and did post analysis by VariScan 2.0^38, 39^ as well as SweepFineder2^40^. For each gene, we used the median of population genetics test statistics as the corresponding value.

### The building of CoVdb

The web interface of CoVdb is on the basis of SWAV^41^. CoVdb also incorporates MSAViewer^42^ to display multiple alignments and phylotree.js^43^ to show the phylogenetic trees. To fit the requirement to display virus data, we made changes in these two softwares, such as changing parameters to fit virus’s dense gene arrangements and adding links within diagrams. The search engine is written by PHP integrated with SQL, BLAT^44^ and NCBI BLAST^45^.

## Data availability

All CoVdb data are publicly and freely accessible at http://covdb.popgenetics.net. Feedback on any aspect of the CoVdb and discussions of Conoravirus gene annotations are welcome by email to.

## Author contributions

Z.Z. developed the web interface of the database, collected and compiled the data, K.M. performed the analysis. Z.Z. and G.M. conceived the idea, coordinated the project and wrote the manuscript.

## Acknowledgments

This work was supported by grants from the National Key Research and Development Program (2019YFC1604600), the National Natural Science Foundation of China (31200941), the Fundamental Research Funds for the Central Universities (106112016CDJXY290002), the National Natural Science Foundation of HeBei province (19226631D).

