## Supplementary Figures for "A database resource for Genome-wide dynamics analysis of Coronaviruses on a historical and global scale"

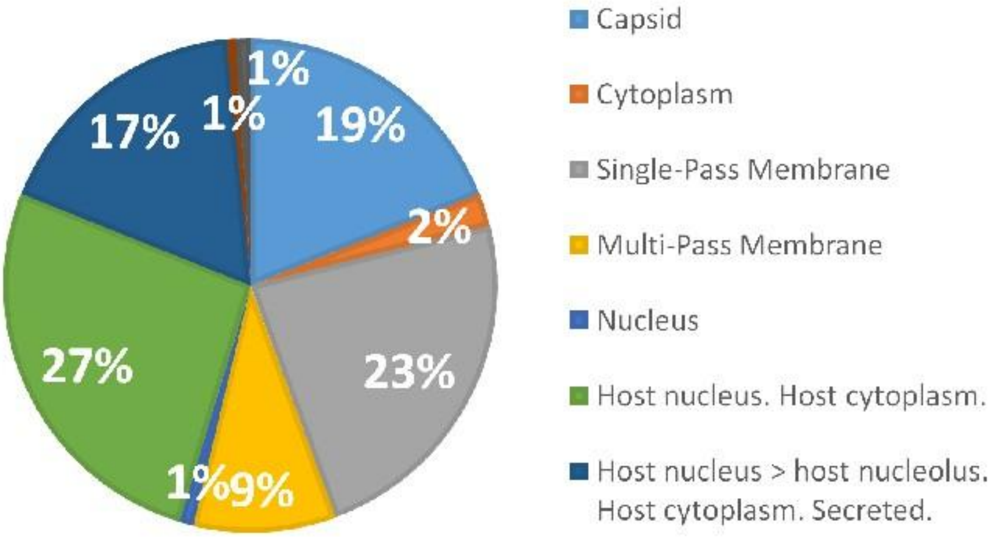


Figure S1. The percentages of Conoravirus gene clusters in each subcellular location group.

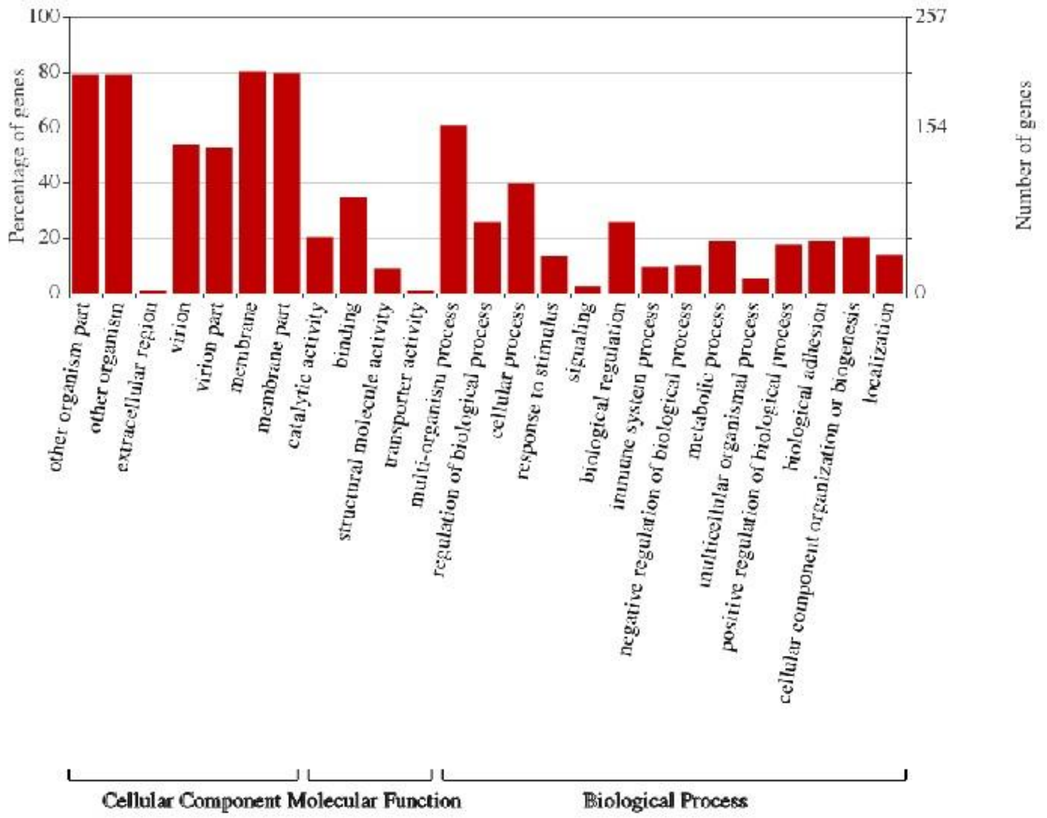


Figure S2. The enrichment of the gene ontology (GO) of Conoravirus gene clusters. This figure was drawn by WEGO.

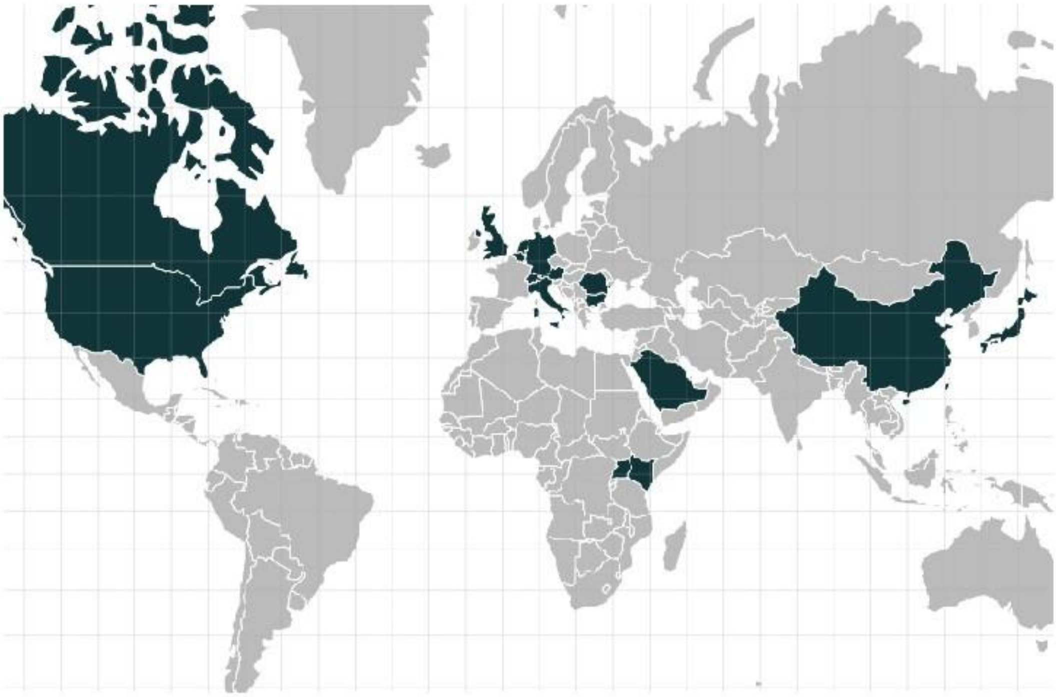

Figure S3. The distribution of strains (marked in dark color) in CoVdb according to country.

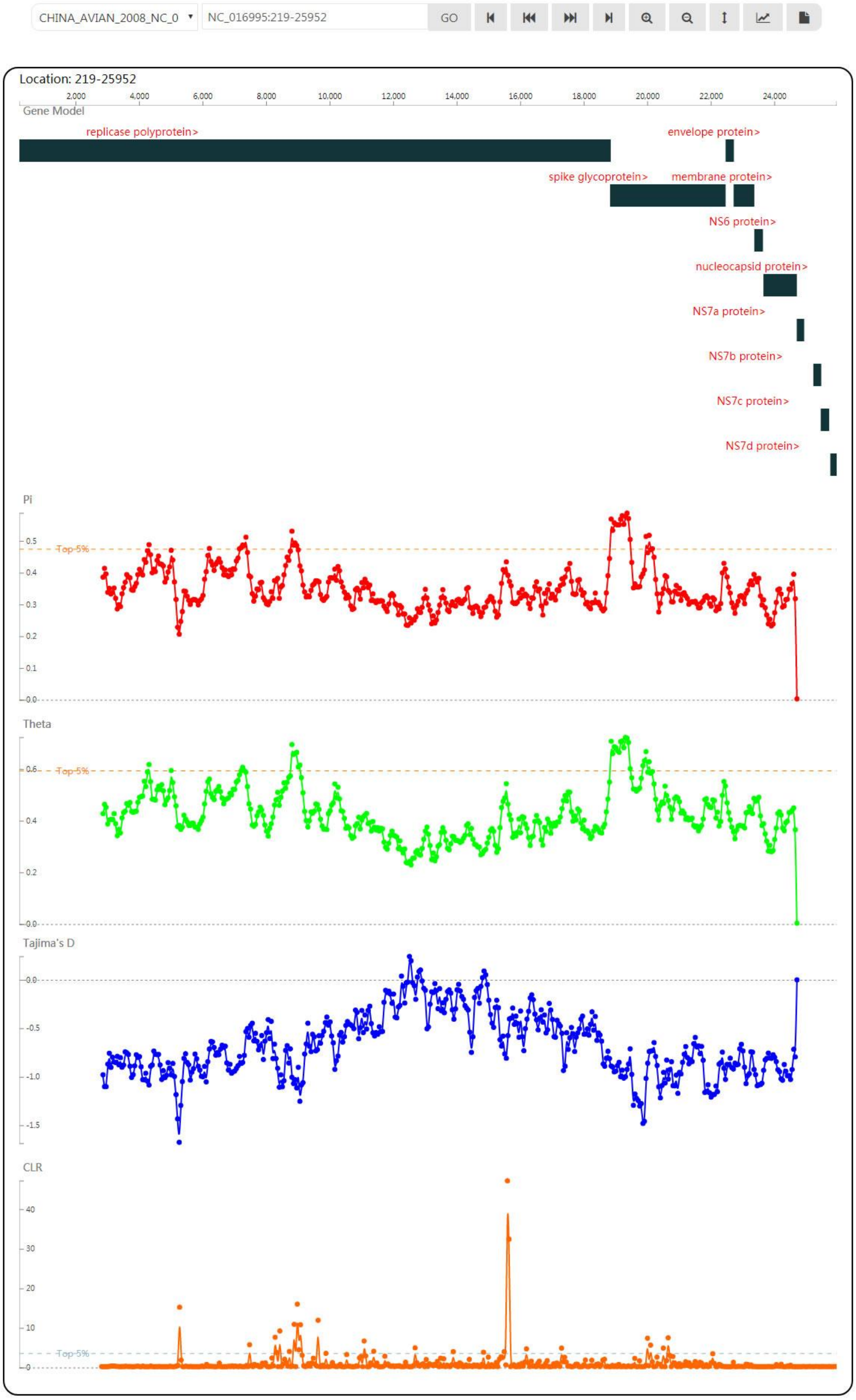


Figure S4. The phylogenetic tree of Coronavirus strains in CoVdb.
